# A novel role for Oxaloacetate Decarboxylase FAHD1 in cardiomyocyte maturation

**DOI:** 10.64898/2026.08.14.744855

**Authors:** Elia Cappuccio, Athanasios Seretis, Attila Kiss, Theresa Zenleser, Max Holzknecht, Donna Paznar, Adolf M. Sandbichler, Christopher Dostal, Maria Cavinato, Jochen Pöling, Bruno K. Podesser, Lisa Schlicker, Almut Schulze, Thomas Braun, Alexander K. H. Weiss, Pidder Jansen-Dürr

## Abstract

Mitochondrial metabolism undergoes dramatic reprogramming during postnatal cardiac maturation, yet the enzymatic regulators that ensure continuity of TCA cycle flux in this period remain incompletely defined. FAHD1 is a mitochondrial oxaloacetate decarboxylase (ODx) with proposed roles in modulating the activity of Complex II of the electron transport chain (ETC), but its physiological relevance *in vivo* has remained unclear. Here, we identify FAHD1 as a critical regulator of mitochondrial function with strong impact on cardiomyocyte (CM) maturation. Using a germline Fahd1-knockout (KO) mouse model, we show that Fahd1 deficiency impairs Complex II respiration, reduces pyruvate levels, and induces a compensatory metabolic shift toward glycolysis and anabolic biosynthesis. Loss of FAHD1 disrupts sarcomere organization, delays the fetal-to-adult myosin isoform switch, and leads to left ventricle systolic dysfunction and cardiomyocyte hypertrophy. These findings highlight FAHD1 as a mitochondrial gatekeeper and potential target for modulating cardiac development and disease.

## 1. Introduction

Mammalian heart development involves a coordinated transition from a fetal to an adult phenotype, encompassing structural maturation of cardiomyocytes (CM) and a metabolic shift from glycolysis to oxidative phosphorylation (OXPHOS)^1^. In mice, this maturation occurs across three phases, embryonic (before P0), neonatal (P0–P30), and juvenile/adult (after P30), with birth triggering a critical transition from hypoxia to normoxia as neonates begin independent respiration^2^.

Shortly after birth, CM exit the cell cycle, become binucleated, and shift from hyperplastic to hypertrophic growth^3^. Concomitantly, the heart switches from carbohydrate-based metabolism to fatty acid oxidation (FAO), which supplies over 95% of ATP in the adult myocardium. This maturation involves not only enhanced oxidative capacity but also changes in the ECM, in the sarcomere organization, and the formation of energetic microdomains to allow maximal mitochondrial ATP production^4^, supporting the dramatic rise in cardiac workload.

Despite extensive studies on embryonic and adult CM regulation, the molecular control of postnatal CM maturation remains incompletely understood^3^. Multi-omics analyses of murine hearts during early postnatal development demonstrated dynamic reprogramming of fatty acid metabolism, ketogenesis, and branched-chain amino acid catabolism^5^, while single-cell RNA sequencing in the heart of WT mice at different age revealed the appearance of discrete developmental trajectories involving either fibroblast-mediated CM maturation or maintaining the proliferative capacity of CM^6^.

Energetic remodeling in the neonatal heart is tightly linked to morphological development. Contractile function requires precise ATP buffering near high-demand sites, mediated by creatine kinase and mitochondrial-anchored phosphotransfer systems. This spatial energetic coupling is enabled by cytoskeletal adaptations and mitochondrial network formation^4^. Structural maturation also includes sarcomere alignment, myosin heavy chain isoform switching (Myh7 to Myh6), and titin isoform transitions, which enhance contractility^7,8^. Whereas it was shown that retinoid X receptor α (RXRα) regulates mitochondrial fatty acid homeostasis (mtFAH) genes essential for perinatal heart function^9^, transcriptional and epigenetic mechanisms integrating structural and metabolic changes in developing CM remain largely undefined.

Fumarylacetoacetate hydrolase domain-containing protein 1 (FAHD1) is the first known eukaryotic oxaloacetate (OAA) decarboxylase^10^. FAHD1 converts OAA to pyruvate via CO₂ release, effectively antagonizing the activity of the anaplerotic enzyme pyruvate carboxylase^11–13^. This reaction regulates mitochondrial OAA levels, which, when elevated, inhibit succinate dehydrogenase (SDH) and impair electron transport by Complex II^14^. Fahd1 deficiency has been shown to induce mitochondrial dysfunction and senescence in human cells^15^, and to impair Complex II activity in cancer cells^16^. Our prior work supports a model wherein FAHD1 functions to maintain Complex II activity by preventing OAA accumulation^17^.

Here, we identify a physiological role for FAHD1 in postnatal heart development. We show that FAHD1 expression peaks during discrete postnatal windows, notably at P3, coinciding with well-established structural and functional transitions in the developing heart. Accordingly, we found that in two-month-old mice, FAHD1 is required for correct sarcomere organization and myosin isoform switching, resulting in proper CM maturation. Of note, FAHD1 deficiency leads to a shift from OXPHOS to glycolysis, accompanied by delayed and altered metabolite profiles, finally resulting in systolic dysfunction and cardiac hypertrophy. Together, these findings suggest that FAHD1 contributes to the structural and metabolic maturation of CM during postnatal development.

## 2. Results

### 2.1. Postnatal FAHD1 expression peaks during critical windows of cardiomyocyte maturation

To define the temporal dynamics of FAHD1 expression during heart development, we analyzed RNA expression data from the Evo-Devo database, which provides cross-species developmental gene expression profiles^18^. In mouse hearts, FAHD1 expression was low during embryogenesis but markedly upregulated during the early postnatal phase, coinciding with the expression pattern of mitochondrial fatty acid homeostasis (mtFAH) signature genes^9^, consistent with a potential role of FAHD1 in postnatal metabolic remodeling.

To assess the FAHD1 expression pattern in cardiomyocytes at the protein level, CM were isolated from WT and Fahd1-KO pups at postnatal days P0, P3, P7, P14, and P21, and FAHD1 protein levels were analyzed by Western blot. FAHD1 expression followed a biphasic pattern, with peak levels at P3 and P21, and reduced expression at intermediate time points (P0, P7, and P14) (**Fig. 1A, B**). These distinct peaks suggest that FAHD1 is required during specific developmental windows, particularly in the first week after birth, a time window known to be associated with rapid metabolic and structural transitions^3,5^.

**Figure 1:**
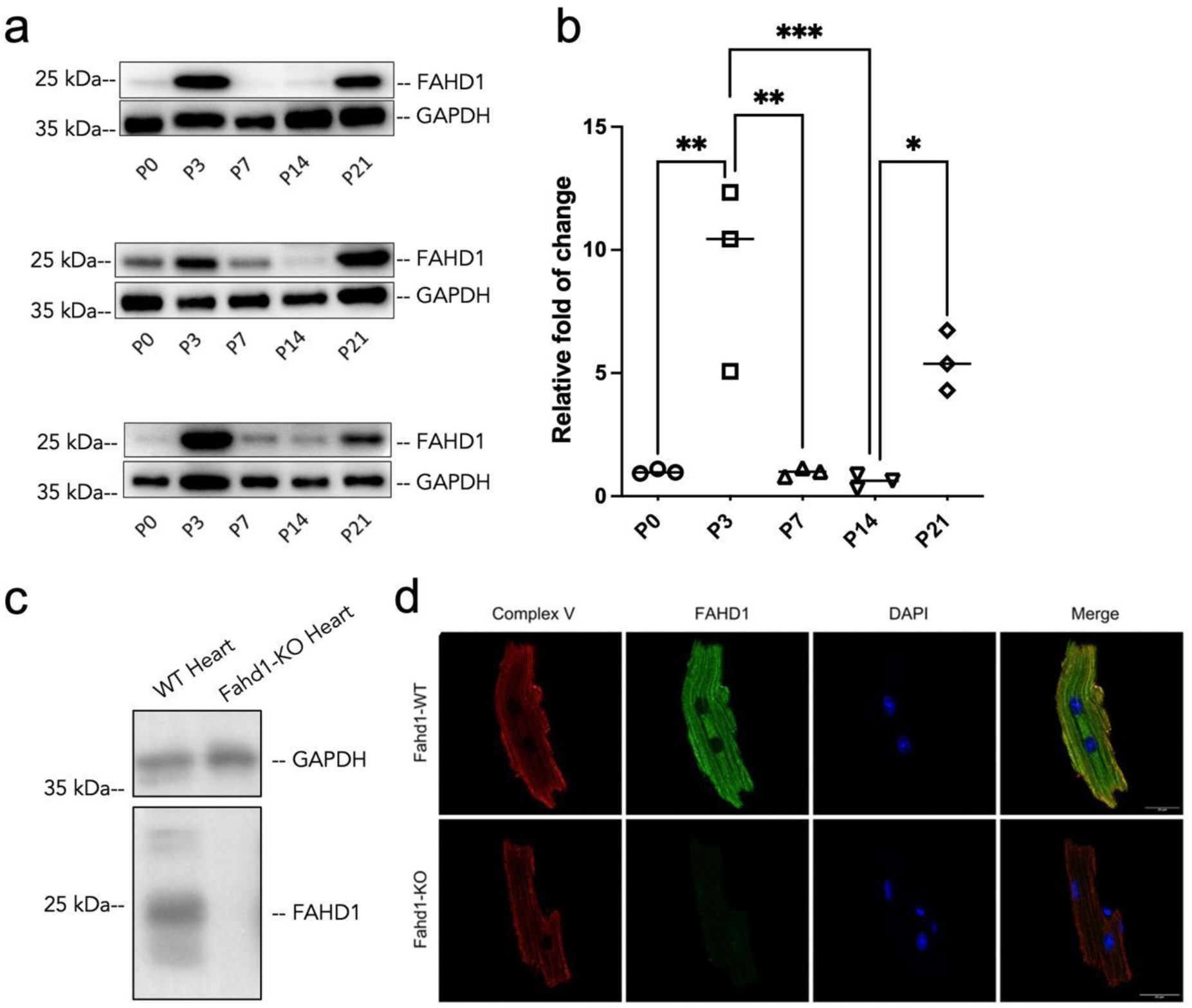
FAHD1 is expressed by cardiomyocytes at different developmental stages: (**A**) Western blot analysis of FAHD1 expression in cardiomyocytes isolated from WT mice at postnatal days P0, P3, P7, P14, and P21. FAHD1 levels display a biphasic temporal pattern, with expression peaks at P3 and P21, corresponding to critical windows of metabolic and structural remodeling. (**B**) Densitometric quantification of FAHD1 protein expression from panel (n=3). Data are presented as median; p-value was calculated with Ordinary one-way ANOVA: *p < 0.05; **p < 0.01; ***p < 0.001. These data highlight the tightly regulated developmental expression of FAHD1 during cardiomyocyte maturation. (**C**) Western blot analysis confirms robust FAHD1 expression in WT hearts of 2 MO mice, with complete absence in Fahd1-KO hearts. (**D**) Immunofluorescence co-staining of FAHD1 (green) and mitochondrial Complex V (red) in WT cardiomyocytes shows colocalization of FAHD1 with mitochondria. Nuclei are counterstained with DAPI (blue). Scale bars: 20 μm.

Western blot analysis confirmed robust FAHD1 expression in WT but not Fahd1-KO hearts **(Figure 1C)**. Immunofluorescence staining further revealed mitochondrial localization of FAHD1 in WT cardiomyocytes, with clear co-localization of FAHD1 with mitochondrial markers (**Figure 1D**), in agreement with previous findings in human cells^15^.

### 2.2. Loss of FAHD1 impairs mitochondrial respiration with compensatory upregulation of glycolysis in cardiomyocytes

To investigate the cellular basis for metabolic consequences of Fahd1 deletion, we examined mitochondrial respiration in isolated cardiomyocytes using high-resolution respirometry (HRR). Fahd1-deficient CM exhibited significantly reduced oxygen consumption following succinate stimulation, indicative of impaired Complex II activity (**Fig. 2A, B**). This finding is consistent with prior reports of Complex II deficiency in FAHD1-depleted cancer cells^16^ and supports our hypothesis that FAHD1 regulates mitochondrial TCA cycle flux by preventing oxaloacetate (OAA)-mediated inhibition of succinate dehydrogenase (SDH).

**Figure 2:**
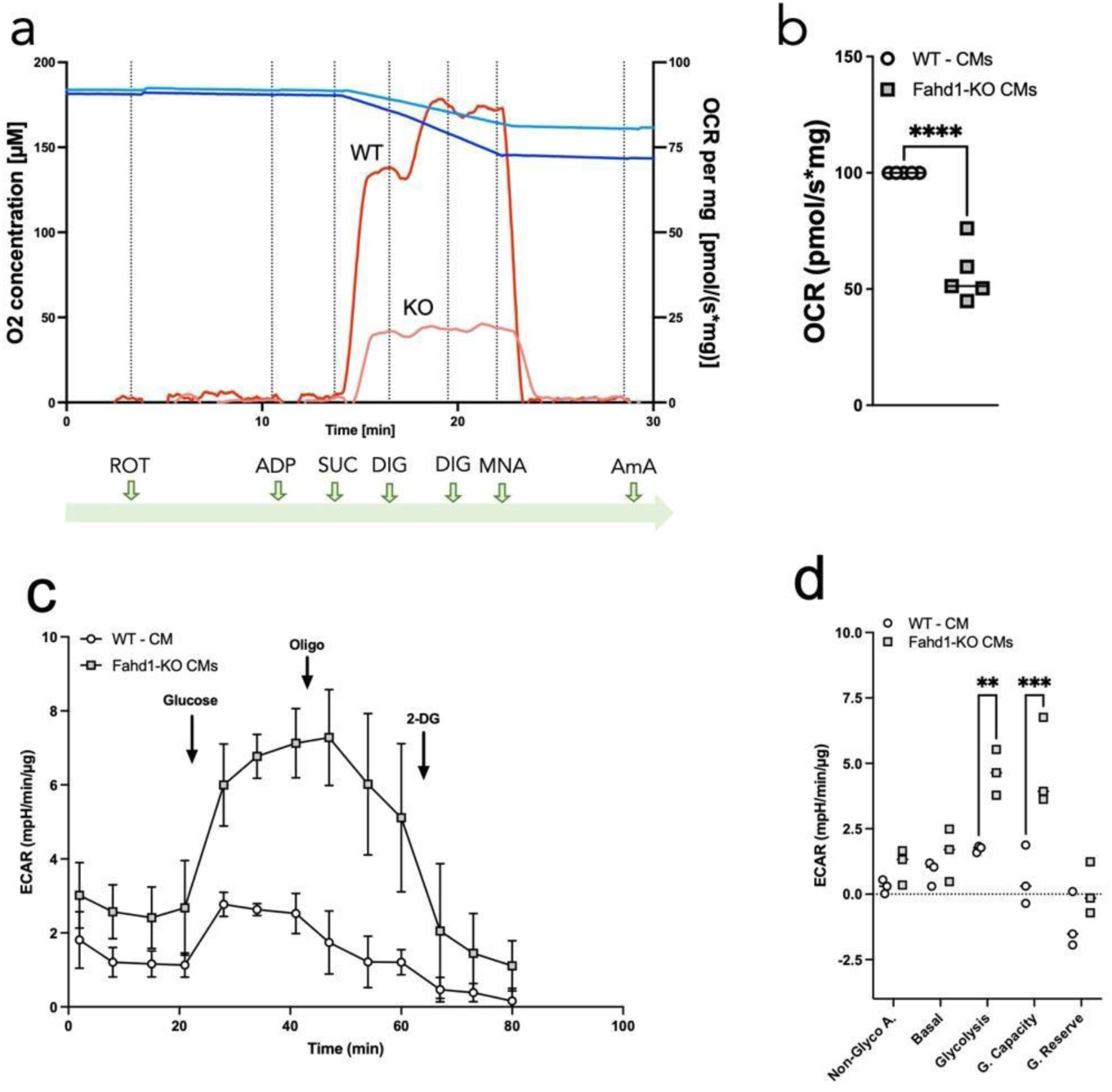
FAHD1 deficiency impairs mitochondrial respiration and induces metabolic reprogramming in cardiomyocytes: (**A**) High-resolution respirometry analysis of mitochondrial Complex II activity in freshly isolated WT and Fahd1-KO cardiomyocytes using the Oroboros Oxygraph O2k system. Digitonin-permeabilized cardiomyocytes were sequentially exposed to ADP, succinate (Complex II substrate), and inhibitors of the electron transport chain (rotenone, malonic acid, antimycin A), as indicated in the timeline. Representative OCR (oxygen consumption rate) traces illustrate reduced Complex II–driven respiration in Fahd1-KO cells. (**B**) Quantification of complex–II-dependent respiration from experiments shown in (A), expressed as a percentage relative to WT controls (set to 100%). Fahd1-KO cardiomyocytes exhibit significantly reduced Complex II activity (n = 5). Data are presented as median; p-value was calculated with unpaired t-test: ****p < 0.0001. (**C**) Glycolytic stress test in WT and Fahd1-KO cardiomyocytes using Seahorse XF analysis (n=3). ECAR (extracellular acidification rate) traces show response to sequential addition of glucose (10 mM), oligomycin (2 mM), and 2-deoxyglucose (50 mM). Fahd1-KO cells display significantly elevated glycolytic capacity relative to WT. All values were normalized to µg of protein. (**D**) Quantification of Glycolysis stress test (n=3), data are presented as median; p-values were calculated with two-way ANOVA, Šídák’s multiple comparisons test: **p < 0.01; ***p < 0.001.

To assess metabolic adaptation to mitochondrial dysfunction, we measured extracellular acidification rates (ECAR) using a glycolytic stress test. Fahd1-KO CM displayed significantly increased glycolytic activity compared to WT controls (**Fig. 2C, D**), indicating a compensatory shift from OXPHOS to glycolysis.

### 2.3. FAHD1 loss promotes a fetal-like metabolic state through altered TCA cycle flux and anabolic metabolism

To further explore the metabolic consequences of FAHD1 deficiency, we performed targeted metabolomics in hearts from two-month-old WT and Fahd1-KO mice. The most prominent alteration was a significant depletion of pyruvate in Fahd1-KO hearts (**Fig. 3**), consistent with the loss of FAHD1, which in WT mice converts oxaloacetate (OAA) to pyruvate^10^. In parallel, we observed an accumulation of nucleotide precursors, including cytidine, adenosine, uridine, and inosine, suggesting a diversion of TCA cycle intermediates toward anabolic biosynthesis, e.g., via the pentose phosphate pathway. These findings suggest that FAHD1 loss initiates a coordinated reversion to a fetal-like metabolic state, characterized by enhanced glycolysis, reduced OXPHOS, and biosynthetic reprogramming. Strikingly, we observed an unexpected imbalance in the concentration of Riboflavin, also known as vitamin B2, which is highly enriched in the heart tissue of Fahd1-KO mice, suggesting that processing of Riboflavin, a well-established precursor for FAD and FMN synthesis, is partially impaired in Fahd1-KO mice. In addition, we also observed a trend for accumulation of 4-hydroxyproline in heart tissue of Fahd1-KO mice, which has been shown to increase levels of HIF-1α presumably by inhibiting its degradation^19^, consistent with the upregulation of glycolysis in Fahd1-KO CM (Fig. 2).

**Figure 3:**
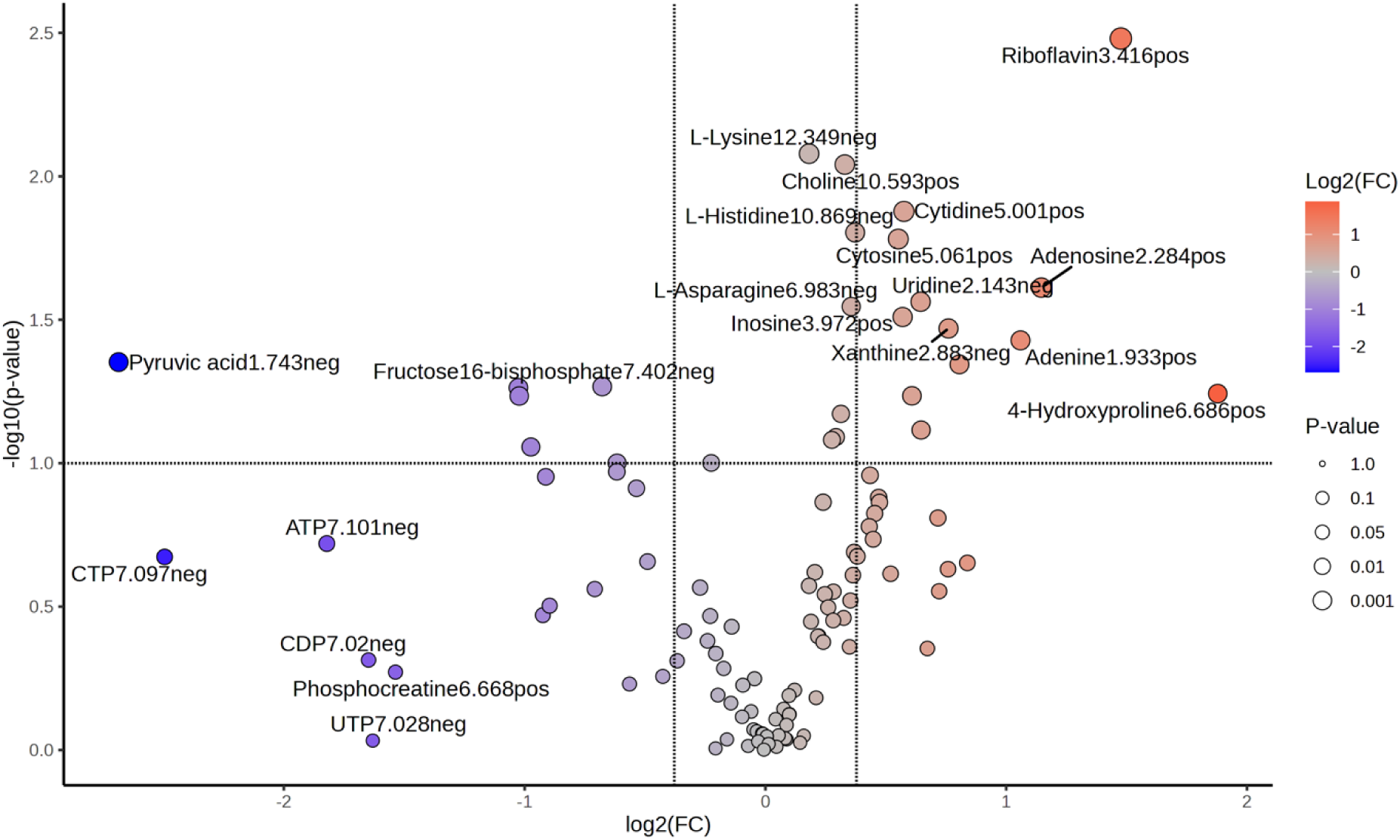
FAHD1 deficiency leads to altered metabolite levels in heart tissue: Targeted metabolomics analysis of hearts from two-month-old WT and Fahd1-KO mice (n=4). The Volcano plot displays all detected metabolites with the 20 most significantly changed metabolites being labeled (name, retention time and mode of detection). Metabolites upregulated in Fahd1-KO heart are shown in red; downregulated metabolites are shown in blue. Statistical analysis was performed using MetaboAnalyst on log2-transformed data. Significance was calculated using unpaired t-tests without FDR correction [p-value threshold 0.1, fold change (FC) threshold 1.3]. Pyruvate depletion is consistent with loss of FAHD1-mediated oxaloacetate decarboxylation.

### 2.4. FAHD1 is required for correct sarcomere organization and contractile protein maturation

Given the mitochondrial dysfunction and metabolic immaturity observed in Fahd1-KO CM, we next investigated whether FAHD1 deficiency also affects structural maturation. Immunofluorescence analysis of α-actinin and F-actin revealed pronounced sarcomeric disorganization in Fahd1-KO CM, in contrast to the highly ordered striated pattern observed in WT cells (**Fig. 4 A**). Quantification of sarcomere length confirmed a significant reduction in Fahd1-KO cells, indicative of an immature contractile apparatus (**Fig. 4 B**). To assess molecular markers of contractile maturation, we analyzed the developmental isoform switch of myosin heavy chains, a well-established indicator of cardiac maturation^7^. In WT hearts, expression of the fetal isoform Myh7 decreased while adult isoform Myh6 increased, consistent with the canonical postnatal transition^7^. In Fahd1-KO hearts, however, this transition was significantly delayed, with persistent Myh7 expression and reduced Myh6 levels (**Fig. 4 C, D**). The decreased Myh6/Myh7 ratio in Fahd1-KO mice suggests a failure to complete the molecular maturation program of sarcomeric proteins.

**Figure 4:**
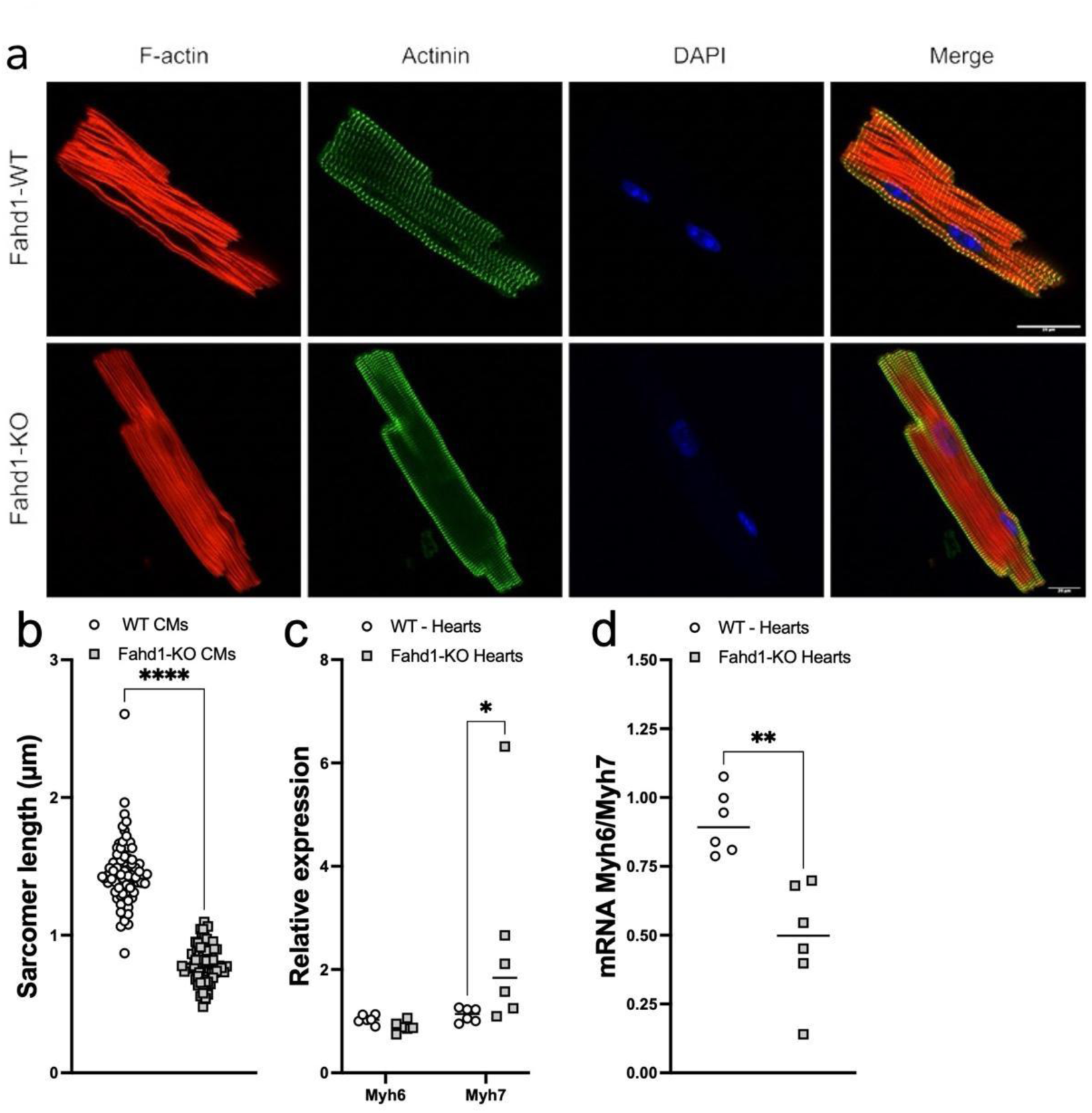
FAHD1 deficiency impairs cardiomyocyte structural maturation and exhibits biphasic developmental expression: (**A**) Immunofluorescence analysis of sarcomere organization in isolated cardiomyocytes from two-month-old WT and Fahd1-KO mice. Sarcomeres were visualized by co-staining for α-actinin (green, Z-discs) and F-actin (red, thin filaments; phalloidin). Fahd1-KO cardiomyocytes display a disorganized filament structure and disrupted sarcomeric alignment compared to the highly ordered striated pattern in WT cells. Scale bars indicate magnification. (**B**) Quantification of sarcomere length, measured as the distance between adjacent α-actinin bands. Fahd1-KO cardiomyocytes show significantly reduced sarcomere length, consistent with an immature contractile phenotype (n = 3). Data are presented as median; p-value was calculated using an unpaired t-test: ****p < 0.0001. (**C**) Quantitative RT-PCR analysis of myosin heavy chain isoforms in hearts from two-month-old WT and Fahd1-KO mice. Fahd1-KO hearts show reduced expression of adult Myh6 (α-MHC) and persistent expression of fetal Myh7 (β-MHC), indicating delayed isoform switching (n = 6). Data are presented as median; p-value was calculated using two-way ANOVA: *p < 0.05 (**D**) Myh6/Myh7 expression ratio calculated from panel (C). Fahd1-KO hearts exhibit a significantly lower ratio, reflecting impaired maturation of the contractile gene program (n = 6). Data are presented as median; p-value was calculated using unpaired t-test: **p < 0.01.

These findings indicate that FAHD1 is not only critical for metabolic maturation but also required for structural remodeling of CM. The persistence of fetal-like sarcomere architecture and contractile protein expression in Fahd1-KO hearts points to a coordinated regulatory role for FAHD1 in aligning energy metabolism with contractile maturation during early postnatal development.

### 2.5. FAHD1 deficiency impairs cardiac function and induces left ventricular hypertrophy

To determine whether FAHD1 plays a physiological role in cardiac function, we performed transthoracic echocardiography in two-month-old Fahd1-KO mice and WT littermates. Fahd1-KO animals exhibited significant systolic dysfunction, as evidenced by a marked reduction in left ventricular ejection fraction (LVEF) compared to WT controls (**Fig. 5A**). This impairment in contractile performance was accompanied by increased left ventricular internal dilatation (LVID) and elevated left ventricular volume (LVV) (**Fig. 5B, C**), indicating compensatory ventricular dilatation.

**Figure 5:**
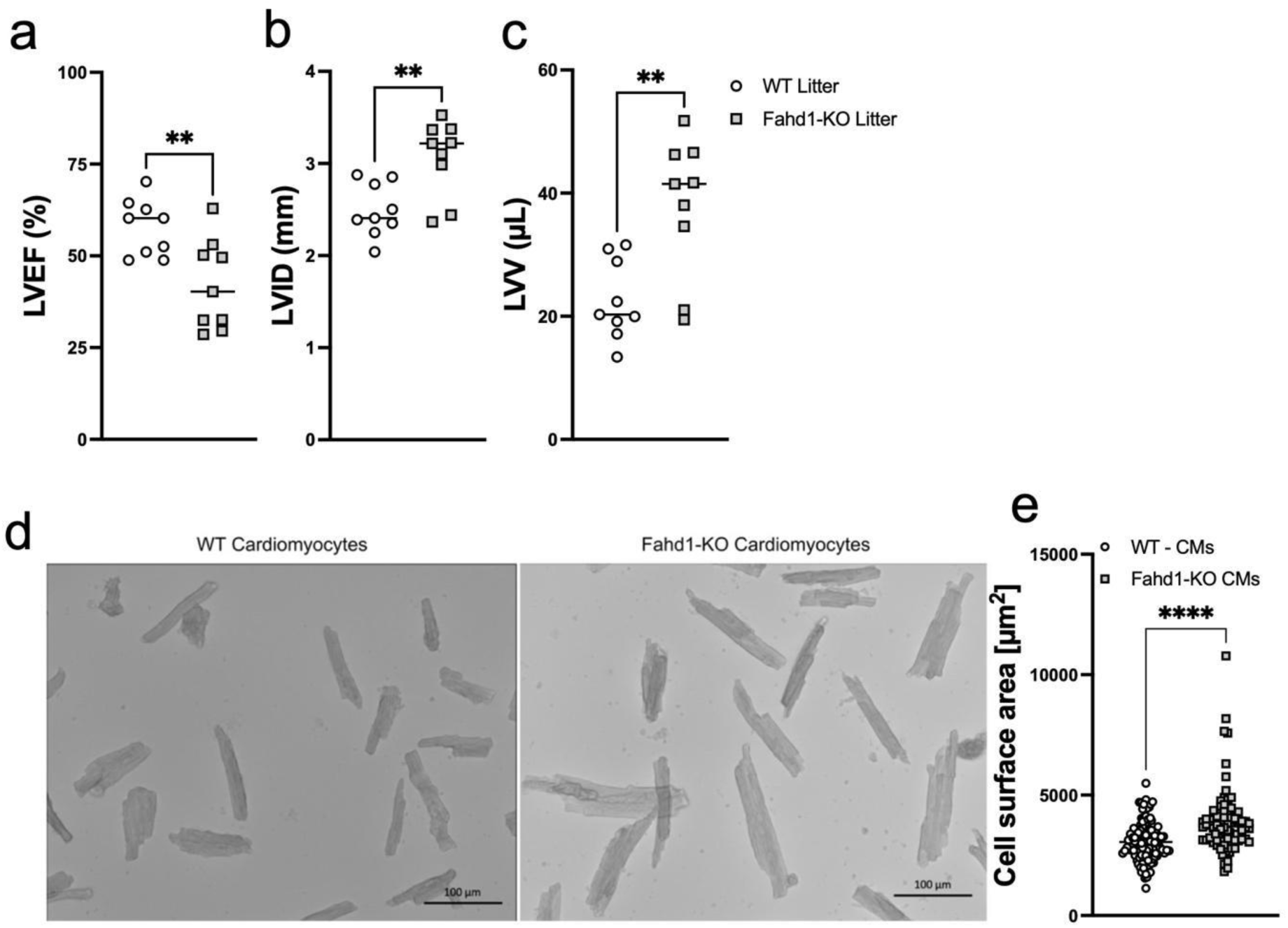
FAHD1 deficiency leads to impaired CM function. (**A–C**) Echocardiographic assessment of cardiac function in two-month-old WT and Fahd1-KO mice (n = 9). (**A**) Left ventricular ejection fraction (LVEF) is significantly reduced in Fahd1-KO mice. (**B**) Left ventricular internal dilatation (LVID) and (**C**) left ventricular volume (LVV) are significantly increased, indicative of compensatory dilation. Data are presented as median; p values calculated with unpaired t-test: **p < 0.01. (**D, E**) Morphometric analysis of isolated cardiomyocytes from two-month-old hearts. (**D**) Representative bright field images show enlarged cardiomyocytes in Fahd1-KO mice relative to WT. (**E**) Quantification of cardiomyocyte surface area reveals significant hypertrophy in Fahd1-KO cells (n = 3). Data are presented as median; p values were calculated with unpaired t test: ****p < 0.0001.

To assess whether these structural changes were reflected at the cellular level, we isolated CM from WT and Fahd1-KO mice for morphometric analysis. Fahd1-KO CM were significantly enlarged, consistent with mild hypertrophy (**Fig. 5D, E**). These findings indicate that FAHD1 is necessary for maintaining normal CM size and cardiac architecture in adulthood.

Together, these data demonstrate that FAHD1 deficiency results in compromised cardiac function and structural remodeling, establishing a previously unrecognized role for FAHD1 in preserving adult myocardial performance.

## 3. Discussion

This study identifies FAHD1 as a novel regulator of CM maturation, acting through its mitochondrial oxaloacetate decarboxylase (ODx) activity to maintain TCA cycle flux and OXPHOS activity in CM, thereby supporting structural and metabolic remodeling during postnatal heart development. Loss of FAHD1 leads to impairment of mitochondrial respiration at the level of Complex II with concomitant upregulation of glycolysis, altered metabolite flux, and persistent features of immature cardiomyocytes, including sarcomeric disorganization and failure to switch to the adult myosin isoform.

The temporal correlation between FAHD1 expression, exhibiting a biphasic pattern with peaks at P3 and P21, and key maturation events supports the hypothesis that FAHD1 contributes to orchestrating postnatal cardiomyocyte development. Given the emerging role of mitochondria in signaling and epigenetic regulation^20^ and the influence of TCA cycle metabolites on chromatin-modifying enzymes^21^, FAHD1 may act upstream of transcriptional and/or epigenetic networks that coordinate the metabolic and contractile maturation of cardiomyocytes. Thus, it is plausible that altered mitochondrial metabolism in the absence of FAHD1 contributes to transcriptional immaturity *via* metabolite– epigenetic crosstalk, a hypothesis warranting further study. Recent studies implicate TCA cycle intermediates as regulators of CM fate and maturation. For example, blockade of FAO through carnitine palmitoyltransferase 1b (Cpt1b) inactivation promotes CM proliferation *via* α-ketoglutarate accumulation and lysine demethylase 5 KDM5 activation^22^. These findings underscore the importance of metabolic flux through the TCA cycle in governing developmental cell fate transitions in the heart.

Metabolic and structural transitions from fetal to adult CM have profound implications for cardiac disease and regeneration. While adult CM show minimal proliferative capacity, fetal CM retain regenerative potential. Reactivation of immature CM phenotypes, via expression of Yamanaka factors (Oct4, Sox2, Klf4, c-Myc), confers regenerative ability in adult hearts^23^. In addition, pathological conditions such as heart failure are associated with reversion to glycolytic metabolism and the reactivation of fetal gene expression programs^24,25^.

Our data suggest that FAHD1 may serve as a significant link between metabolic and structural maturation. At the structural level, FAHD1 deficiency delays key maturation events, including sarcomere alignment and Myh6 induction. The sustained expression of fetal Myh7, shorter sarcomeres, and disorganized cytoskeletal architecture in Fahd1-KO cardiomyocytes are consistent with a failure to fully engage the adult contractile program.

Mapping metabolite changes onto cardiac metabolic pathways revealed, in addition to the shift from OXPHOS to glycolysis (**Fig. 2**), a potential shift toward the pentose phosphate pathway (upregulation of the nucleotides adenosine, cytidine, inosine, and uridine), reminiscent of fetal cardiomyocytes (**Fig. 6**). Reprogramming of the metabolism in Fahd1-KO hearts likely serves to support increased nucleotide and amino acid synthesis in response to impaired mitochondrial energy production. The observed metabolite profile aligns with mechanisms previously described in regenerative CM states and supports the idea that mitochondrial TCA cycle flux is a critical determinant of cardiomyocyte maturation and identity^26^. Decreased levels of pyruvate in Fahd1-KO mice probably reflect the reduced decarboxylation of OAA in the absence of FAHD1, as FAHD1 has been shown to decarboxylate OAA in vitro and in vivo^8^. On the other hand, it is conceivable that enhanced conversion of pyruvate to lactate, evidenced by an increased extracellular acidification rate (**Fig. 2**), also contributes to the observed depletion of pyruvate.

**Figure 6:**
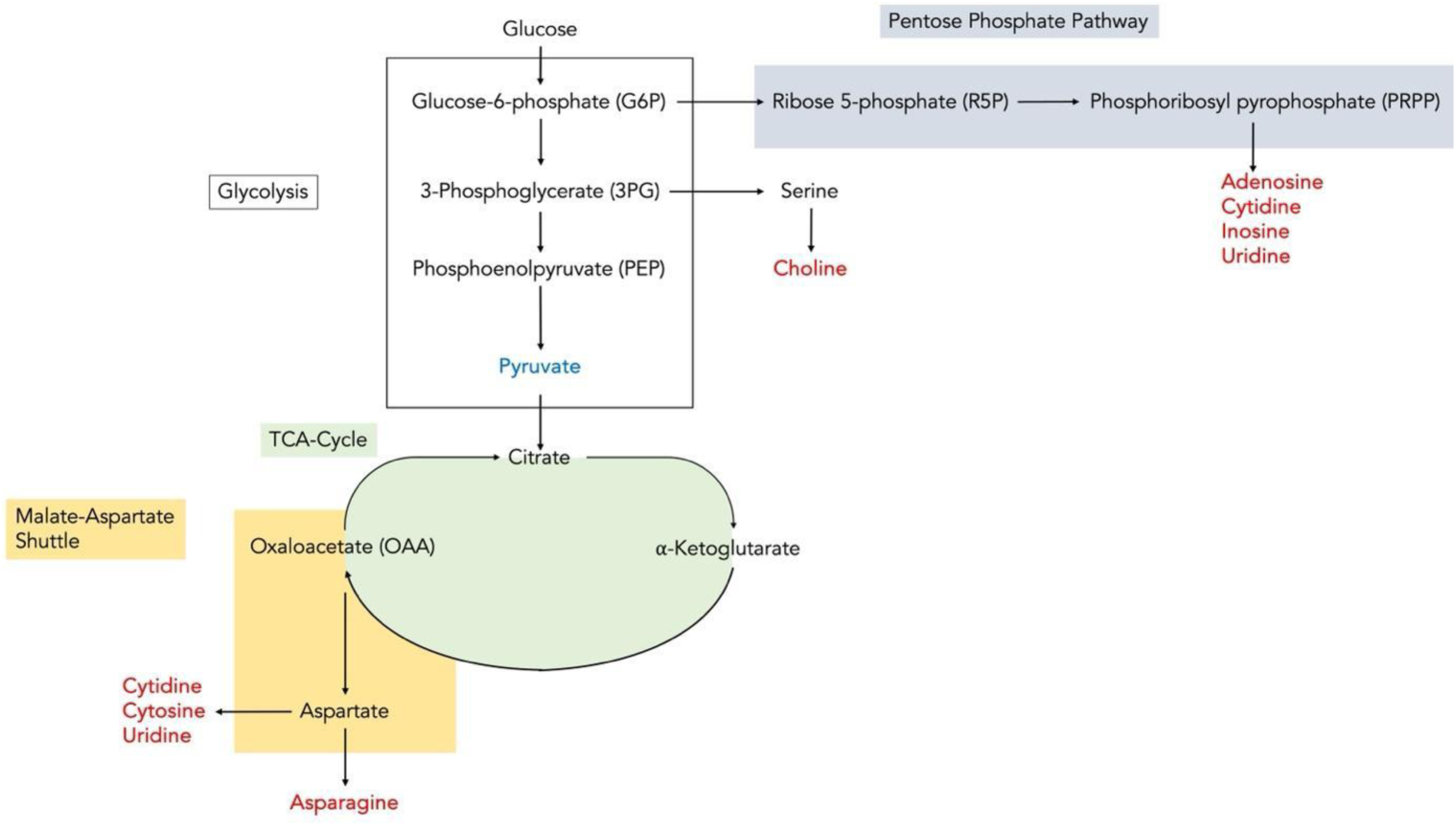
FAHD1 deficiency leads to metabolic rewiring in the mouse heart: Schematic of metabolic alterations in Fahd1-KO hearts based on data in Figure 3. The metabolite profile indicates a shift toward the pentose phosphate pathway to support anabolic metabolism. Concurrent activation of the malate–aspartate shuttle suggests a compensatory mechanism to remove mitochondrial OAA and partially preserve Complex II function in the absence of FAHD1.

The elevated levels of pyrimidine nucleotide precursors cytidine, cytosine and uridine, as well as the amino acid asparagine, all requiring cytosolic aspartate for their synthesis, may be indicative of the activation of the malate-aspartate shuttle (**Fig. 6**), well known as an alternative OAA-consuming pathway, in the hearts of Fahd1-KO mice, which could allow to compensate for the observed Complex II defect in Fahd1-KO CM. In addition, the partial recovery of Complex II activity observed in Fahd1-KO CM of two-month-old Fahd1-KO mice may reflect the compensatory activation of PCK2, representing yet another alternative OAA-consuming pathway^13^. More work will be required to delineate the exact metabolic rewiring caused by FAHD1 deficiency in cardiomyocytes.

Another intriguing finding is the increased concentration of Riboflavin (Vitamin B2) in heart tissue of Fahd1-KO mice. Since mammalian cells cannot synthesize Riboflavin, it appears that conversion of Riboflavin to important metabolites, such as Flavin adenine dinucleotide (FAD), is reduced in Fahd1-KO hearts. This could indicate that FAHD1-deficient hearts may not be able to convert Riboflavin into FAD. As FAD is a cofactor for many enzymes, including short-chain acyl-CoA dehydrogenase (SCAD) and components of the respiratory chain, like SDH, losing the ability to make this metabolite could have adverse effects. FAD-dependent enzymes influence cardiac health by affecting fatty acid oxidation, energy production, and by protecting against pathological hypertrophy and fibrosis. In this context, we observed a trend toward diminished levels of both ATP and phosphocreatine in heart tissue from Fahd1-KO mice (**Fig. 3**), suggesting that Fahd1-KO hearts experience energy starvation. This energy deficit may also impair FAD synthesis from riboflavin, a process that requires ATP.

In conclusion, our findings establish FAHD1 as a critical regulator of postnatal cardiac maturation through its oxaloacetate decarboxylase activity. By irreversibly converting OAA to pyruvate, FAHD1 sustains TCA cycle flux and prevents OAA-mediated inhibition of Complex II. Loss of this function disrupts mitochondrial respiration, reprograms cardiac metabolism, and impairs structural maturation of cardiomyocytes.

## Limitations of the study

While our study demonstrates a strong link between FAHD1 function and CM maturation, the use of a global Fahd1 knockout model does not exclude potential systemic or developmental compensation. Although we performed detailed cardiomyocyte-specific phenotyping and used isolated primary cells for mechanistic analyses, future studies employing cardiomyocyte-specific FAHD1 deletion (e.g., via Myh6-Cre) will be essential to confirm the cell-autonomous role of FAHD1 in cardiac development *in vivo*. While Fahd1-KO mice remain viable, this observation likely reflects compensatory metabolic adaptations in specific tissues or a developmentally restricted requirement for FAHD1 that has yet to be fully elucidated. Downregulation of Fructose 1,6-bisphosphate, usually a good indicator of glycolysis, in the heart tissue of Fahd1-KO mice (**Fig. 3**) seems to be at odds with the upregulation of glycolytic activity observed in freshly isolated CM from Fahd1-KO mice (**Fig. 2**). Whereas the underlying mechanisms for this phenomenon remain to be clarified, it is possible that it reflects metabolic regulation in other cardiac cell types, distinct from CM.

## Outlook

Notably, the metabolic and structural alterations observed in Fahd1-deficient hearts, such as increased glycolysis, reduced mitochondrial respiration, and persistent fetal gene expression, closely mirror features reported in immature or failing human cardiomyocytes. The resemblance between the Fahd1-KO phenotype and metabolic cardiomyopathies supports further investigation into FAHD1 expression or genetic variation in the context of congenital heart disease, pediatric cardiomyopathy, and heart failure. Moreover, exploring FAHD1 function in human iPSC-derived cardiomyocytes may provide valuable translational insight into cardiac maturation, regenerative strategies, and disease modeling.

## Abbreviations

ASP: Aspartate
αKG: Alpha-ketoglutarate
BCA: Bicinchoninic acid
BDM: 2,3-Butanedione monoxime
CM: Cardiomyocyte
CIT: Citrate
DAPI: 4′,6-Diamidino-2-phenylindole
DMEM: Dulbecco’s Modified Eagle Medium
ECAR: Extracellular acidification rate
ETC: Electron transport chain
FAHD1: Fumarylacetoacetate hydrolase domain-containing protein 1
FUM: Fumarate
GAPDH: Glyceraldehyde 3-phosphate dehydrogenase
GOT2: Glutamic-oxaloacetic transaminase 2
HRR: High-resolution respirometry
KO: Knockout
LC-MS/MS: Liquid chromatography–tandem mass spectrometry
LV: Left ventricle / ventricular
LVEF: Left ventricular ejection fraction
LVID: Left ventricular internal diameter
LVV: Left ventricular volume
MAL: Malate
Myh6: Alpha-myosin heavy chain
Myh7: Beta-myosin heavy chain
OAA: Oxaloacetate
OCR: Oxygen consumption rate
ODx: Oxaloacetate decarboxylase
PBS: Phosphate-buffered saline
PCK2: Phosphoenolpyruvate carboxykinase 2 (mitochondrial)
PEP: Phosphoenolpyruvate
PFA: Paraformaldehyde
PVDF: Polyvinylidene difluoride
qRT-PCR: Quantitative reverse transcription polymerase chain reaction
RIPA: Radioimmunoprecipitation assay
SDH: Succinate dehydrogenase
SEM: Standard error of the mean
SUC: Succinate
TA: Tautomerase activity
TCA: Tricarboxylic acid (cycle)
WT: Wild type
2-DG: 2-Deoxyglucose

## Acknowledgements.

This research was partially funded by the Austrian Science Fund (FWF), Grant-DOI 10.55776/FG24 (SENIOPROM) to Pidder Jansen-Dürr.

## Author Contributions

E.C., A.S., J.P., T.B. and P.J. conceived the study and designed the experiments. E.C., T.Z., D.P., M.C., L.S., and A.S. performed experiments and collected data. A.M.S. contributed to methodology development and provided technical support. A.K.H.W. and A.S. conducted data analysis and interpretation. E.C., A.S., A.K.H.W. and P.J. wrote the manuscript with input from all authors. P.J. supervised the project. All authors reviewed and approved the final manuscript.

## Competing Interest

All authors declare that they have no conflicts of interest.

## MATERIAL AND METHODS

### Animals

Fahd1-knockout (Fahd1-KO) mice were generated as previously described^10^. Mice were housed and bred at the animal facility of the Center of Chemistry and Biomedicine, Innsbruck, under specific-pathogen-free conditions (SPF). All experiments were performed on 2-month-old, male animals. Group allocation was done according to the animals’ genotype. All animal procedures were approved by the institutional ethics committee and conducted in accordance with national and EU regulations for animal experimentation.

## PROCEDURES

### Echocardiography

Cardiac function was assessed in male two-month-old wild-type (WT) and Fahd1-knockout (Fahd1-KO) mice using transthoracic echocardiography as described previously^27^. Mice were anesthetized with 3-4% of isoflurane for induction and maintained at 1-1.5% of isoflurane in oxygen (1.5 L/min) during imaging. The body temperature was recorded and maintained at 37 °C using a heated platform, and electrocardiogram (ECG) signals were recorded via paw electrodes embedded in electrode gel. Images were acquired using a Vevo 3100 high-resolution imaging system (FUJIFILM VisualSonics) equipped with an MX550D transducer. The parasternal short-axis view at the mid-papillary level was used to evaluate left ventricular function. M-mode recordings were analyzed using Vevo®Lab software (Fujifilm VisualSonics).

### Cardiomyocyte Isolation

Cardiomyocytes (CM) were isolated from mice at various developmental stages (E17.5, P0, P3, P7, P14, and P21) using stage-specific protocols adapted from established methods^6,22,28^.

For embryonic and early postnatal stages (E17.5–P3), hearts were excised, washed in phosphate-buffered saline (PBS), and ventricles were minced into 1–3 mm³ fragments. Tissue was plated onto fibronectin-coated dishes and cultured in primary neonatal cardiomyocyte medium [DMEM/M199 supplemented with 10% horse serum, 5% fetal calf serum, and antibiotics]. These cells, which attach and spread on the dish surface, are referred to as *adherent CM*.

For later postnatal stages (P7–P30), hearts were excised, washed in calcium- and magnesium-free PBS containing 20 mM 2,3-butanedione monoxime (BDM), and minced. Tissue was incubated overnight at 4°C in digestion buffer containing HBSS (without Ca²⁺/Mg²⁺), 20 mM BDM, and 0.0125% trypsin. The next day, tissue fragments were enzymatically digested at 37°C using 0.75 mg/mL collagenase/dispase in L15 medium with BDM.

Cell suspensions were filtered through 40–100 µm strainers, centrifuged at 100 × g for 5 minutes, and resuspended in plating medium (65% high-glucose DMEM, 19% M-199, 10% horse serum, 5% fetal calf serum, 1% penicillin/streptomycin). To enrich for cardiomyocytes; fibroblasts and endothelial cells were removed *via* pre-plating: cells were incubated on uncoated 10 cm dishes for 1–3 hours at 37°C, allowing non-CM to adhere. The remaining *non-adherent CM* were collected for downstream analyses.

For CM isolation of adult mice (2 months old), was performed as previously described^29^. In brief, mice were euthanized by isoflurane inhalation, the chest cavity opened, and the vena cava severed. The right ventricle was washed by EDTA buffer injection (NaCl 130 mM, KCl 5 mM, NaH_2_PO_4_ 0.5 mM, HEPES 10 mM, Glucose 10 mM, BDM 10 mM, Taurine 10 mM, and EDTA 5mM). Subsequently, the aorta was clamped, the heart removed and placed into a 60 mm dish and the left ventricle was washed by EDTA buffer injection. Next, the heart was washed by Perfusion buffer (NaCl 130 mM, KCl 5 mM, NaH_2_PO_4_ 0.5 mM, HEPES 10 mM, Glucose 10 mM, BDM 10 mM, Taurine 10 mM, and MgCl_2_ 1mM) injection into the left ventricle. After the second wash the heart was digested by consecutive, intra-ventricular injections of Liberase (0.125 mg/ml in Perfusion buffer). When the tissue was visibly digested, the digestion was stopped by the addition of 10% FBS and the heart tissue was mechanically dissolved into 5 mm pieces, passed through a 100 µm cell strainer and allowed to precipitate by gravity settling. After precipitation, the supernatant was removed and calcium was gradually reintroduced by washing the cell pellet in Perfusion buffer containing increasing concentrations (0.34 mM, 0.68 mM, and 1.02 mM) of calcium and centrifugating the cells at 500 rpm for 10 sec.

### Immunofluorescence Staining and Microscopy

Cardiomyocytes were fixed in 4% paraformaldehyde (PFA) for 20 minutes at room temperature and permeabilized with 0.3% Triton X-100 in PBS for 5 minutes. Cells were blocked in staining buffer (2% BSA) for 20 minutes at room temperature. The cells were incubated with primary antibodies against FAHD1 (1:1000), and Complex V (A21351 - Invitrogen; 1:250), in staining buffer overnight at 4°C. Cells were washed in staining buffer and incubated with species-specific secondary antibodies (1:2000 Goat anti-rabbit Alexa Fluor 488, A27034 - Thermofisher Scientific; 1:500 goat anti-mouse Alexa Fluor 546, A11003-Invitrogen) for 1 hour at room temperature. For α-ACTININ (1:500, 130-106-995 - Miltenyi), after incubation with primary AB, CM were incubated with a mouse α-BIOTIN (1:1000 ab 201341 - Abcam) secondary AB and finally with a goat anti-mouse Alexa Fluor 488 AB (1:1500, A27034 - Thermofisher Scientific). F-actin was stained using fluorescently conjugated phalloidin (1:500, A12380 - Invitrogen), and nuclei were counterstained with DAPI. Fluorescent images were acquired using a Yokogawa CV1000 spinning disk confocal system (Olympus) and a SP5-II confocal laser scanning microscope (Leica). Images were processed using ImageJ (NIH) and deconvolved with Huygens Professional software (Scientific Volume Imaging), following standard protocols^30^. Sarcomere length was quantified as the distance between adjacent α-actinin bands using ImageJ line scan analysis. At least 50 sarcomeres per sample were measured across three biological replicates.

### Western Blotting

Protein extracts were prepared from freshly isolated cardiomyocytes or ventricular heart tissue by homogenization in RIPA buffer (Thermo Fisher Scientific) supplemented with protease and phosphatase inhibitors (Roche). Protein concentrations were determined using the Pierce BCA Protein Assay Kit (Thermo Fisher Scientific). Equal amounts of protein (20–40 µg) were used. Band intensities were quantified using ImageJ software and normalized to GAPDH. All Western blot experiments were repeated in at least three independent biological replicates. Antibodies used: rabbit anti-mouse FAHD1 (1:3000 in 5 % BSA in TBS-T), GAPDH (1:5000 in 5 % BSA in TBS-T; 10494-1-AP - Proteintech).

### Gene Expression Analysis

Total RNA was extracted from snap-frozen left ventricular tissue using TRIzol reagent (Thermo Fisher Scientific) according to the manufacturer’s instructions. Quantitative real-time PCR (qRT-PCR) was performed using SYBR Green Master Mix (Bio-Rad) on a CFX96 Touch Real-Time PCR Detection System (Bio-Rad). Relative expression levels were calculated using the ΔΔCt method, with Gapdh as the endogenous reference gene. All reactions were performed in technical triplicate with at least three biological replicates per genotype.

### Metabolic Flux Analysis

Glycolytic function and mitochondrial stress responses were assessed using a Seahorse XFp Analyzer (Agilent Technologies) to measure extracellular acidification rate (ECAR) and oxygen consumption rate (OCR), respectively. Cardiomyocytes isolated from WT and Fahd1-KO mice were plated at a density of 1,000 cells per well in Seahorse XF culture plates in Seahorse assay medium and were allowed to settle by gravity for 30 minutes in a non-CO₂ incubator at 37°C before analysis. Glycolysis (103017-100, Agilent) and Mito stress test (103010-100, Agilent) were performed according to the manufacturer’s instructions. All data were normalized to total protein content per well, assessed by BCA assay.

### High-Resolution Respirometry (HRR)

Mitochondrial respiratory function was assessed in freshly isolated cardiomyocytes from two-month-old WT and Fahd1-KO mice using high-resolution respirometry (HRR). Oxygen consumption rates (OCR) were measured using an Oroboros Oxygraph-2k system (Oroboros Instruments) at 37°C. Basal respiration was recorded following the addition of substrates including ADP (2.5 mM) and succinate (100 mM), followed by sequential additions of rotenone (0.5 µM), malonic acid (15 mM), and antimycin A (2.5 µM) to isolate Complex II-dependent respiration and inhibit upstream and downstream respiratory chain components. Respiratory flux was normalized to total protein content, determined by BCA. Complex II activity was calculated as the rotenone-sensitive, malonic acid-inhibited OCR in the presence of succinate.

### Metabolomics

Targeted metabolomic profiling was performed on left ventricular tissue isolated from two-month-old wild-type (WT) and Fahd1-knockout (Fahd1-KO) mice. Heart tissue was rapidly excised, snap-frozen in liquid nitrogen, and stored at −80 °C until processing. 15 mg of heart tissue was homogenized in 400 µl ice-cold mixture of methanol/water/acetonitrile (50/20/30, v/v/v) with internal standards (IS: MSK-CAA-1, DLM-7654, DLM-9476, DLM-9045, DLM-9071, DLM-6068, DLM-831, DLM-3487 (Cambridge Isotope Laboratories)) using 1.4 mm ceramic beads in a Retsch mill. Polar metabolites were extracted using STRATA C18-E solid phase columns (8B-S001-DAK, Phenomenex), which were previously activated with acetonitrile and equilibrated with methanol/water/acetonitrile (50/20/30, v/v/v). After loading the homogenized samples, the eluate was dried in a vacuum concentrator (Labconco). Samples were dissolved in 100 μL 5 mM Ammonium Acetate in Acetonitrile/Water (75:25, v/v) prior to LC-MS analysis.

Samples were analyzed using liquid chromatography coupled to high-resolution mass spectrometry (LC-HRMS) on a DIONEX Ultimate 3000 UPLC system coupled to a Q Exactive Plus mass spectrometer (Thermo Fisher). Polar metabolites were separated on an amide-HILIC column (2.6 μm, 2.1x100 mm, Thermo Fisher) using 5 mM Ammonium Acetate in 5% Acetontrile as solvent A and 5 mM Ammonium Acetate in 95% Acetontrile as solvent B. The gradient started with 98% solvent B for 2 min, which was followed by a decrease to 40% B within 5 minutes. After maintaining 40% for 13 minutes, the gradient returned to 98% within 1 min, and the column was re-equilibrated for 5 more minutes. The injection volume was set to 3 µl, and the flow rate was kept at 350 µl/min. The polar metabolites were ionized in a hESI ion source, and the scan range was set to 69.0-1000 m/z with a resolution of 70,000, operated in polarity switching mode (positive and negative ionization). Peaks were integrated using El-MAVEN (v.12.1-beta, Elucidata) by comparing m/z using in-house databases at an accuracy of ±5 ppm. Retention time and fragmentation patterns were used to confirm metabolite identity.

Data were normalized to IS and total protein concentration (measured by BCA assay) and expressed as relative abundance or fold change compared to WT controls. Metabolite heatmaps and pathway maps were generated using MetaboAnalyst^31^. Significantly altered metabolites were defined as those with an absolute fold change ≥1.5 and a p-value <0.05 (unpaired two-tailed t-test).

### Quantification and Statistical Analysis

All data are presented as mean ± standard error of the mean (SEM) unless otherwise indicated. Statistical analyses were performed using GraphPad Prism 10. For comparisons between two groups, unpaired two-tailed Student’s t-tests were used. For multiple group comparisons, two-way ANOVA followed by Sidak’s test was applied, as appropriate. Normality was assessed using the Shapiro-Wilk test, and variance equality was evaluated with an F-test or Levene’s test.

Animal numbers (n) used for each experiment are indicated in the figure legends. A p-value of <0.05 was considered statistically significant. No statistical methods were used to pre-determine sample size, but sample sizes are consistent with standard practice in the field. Experiments were not randomized, and investigators were not blinded to group allocation, except where explicitly stated.

## SUPPLEMENTARY MATERIAL

**Supplementary Table 1**: Relative metabolite concentrations in the heart of 2-month-old Fahd1-KO vs WT mice

## Notes

### Competing Interest Statement

The authors have declared no competing interest.

## References

1. Lopaschuk, G. D. & Jaswal, J. S. Energy metabolic phenotype of the cardiomyocyte during development, differentiation, and postnatal maturation. J Cardiovasc Pharmacol 56, 130–140 (2010).

2. Taliani, V. et al. The long noncoding RNA Charme supervises cardiomyocyte maturation by controlling cell differentiation programs in the developing heart. Elife 12, 1–29 (2023).

3. Velayutham, N., Agnew, E. J. & Yutzey, K. E. Postnatal Cardiac Development and Regenerative Potential in Large Mammals. Pediatr Cardiol 40, 1345–1358 (2019).

4. Piquereau, J. et al. Postnatal development of mouse heart: formation of energetic microdomains. J Physiol 588, 2443–2454 (2010).

5. Talman, V. et al. Molecular Atlas of Postnatal Mouse Heart Development. J Am Heart Assoc 7, (2018).

6. Zhu, M. et al. Distinct mononuclear diploid cardiac subpopulation with minimal cell–cell communications persists in embryonic and adult mammalian heart. Front Med 1–18 (2023) doi:10.1007/s11684-023-0987-9.

7. Yin, Z., Ren, J. & Guo, W. Sarcomeric protein isoform transitions in cardiac muscle: a journey to heart failure. Biochim Biophys Acta 1852, 47–52 (2015).

8. Sanger, J. W. et al. Assembly and maintenance of myofibrils in striated muscle. Handb Exp Pharmacol 235, 39–75 (2017).

9. Paredes, A. et al. γ-Linolenic acid in maternal milk drives cardiac metabolic maturation. Nature 618, 365–373 (2023).

10. Pircher, H. et al. Identification of FAH Domain-containing Protein 1 (FAHD1) as Oxaloacetate Decarboxylase. Journal of Biological Chemistry 290, 6755–6762 (2015).

11. Weiss, A. K. H. et al. Inhibitors of Fumarylacetoacetate Hydrolase Domain Containing Protein 1 (FAHD1). Molecules 2021, Vol. 26, Page 5009 26, 5009 (2021).

12. Zmuda, A. J. et al. A universal metabolite repair enzyme removes a strong inhibitor of the TCA cycle. Nat Commun 15, (2024).

13. Cappuccio, E. et al. FAHD1 and mitochondrial metabolism: a decade of pioneering discoveries. FEBS J 10.1111/febs.17345 (2024) doi:10.1111/febs.17345.

14. Stepanova, A., Shurubor, Y., Valsecchi, F., Manfredi, G. & Galkin, A. Differential susceptibility of mitochondrial complex II to inhibition by oxaloacetate in brain and heart. Biochim Biophys Acta 1857, 1561–1568 (2016).

15. Petit, M., Koziel, R., Etemad, S., Pircher, H. & Jansen-Dürr, P. Depletion of oxaloacetate decarboxylase FAHD1 inhibits mitochondrial electron transport and induces cellular senescence in human endothelial cells. Exp Gerontol 92, 7–12 (2017).

16. Holzknecht, M. et al. The mitochondrial enzyme FAHD1 regulates complex II activity in breast cancer cells and is indispensable for basal BT-20 cells in vitro. FEBS Lett 10.1002/1873-3468.14462 (2022) doi:10.1002/1873-3468.14462.

17. Etemad, S. et al. Oxaloacetate decarboxylase FAHD1 – a new regulator of mitochondrial function and senescence. Mech Ageing Dev 177, 22–29 (2019).

18. Cardoso-Moreira, M. Gene expression across mammalian organ development-manuscript - supplement. Nature 571, 1–5 (2019).

19. Phang, J. M. The regulatory mechanisms of proline and hydroxyproline metabolism: Recent advances in perspective. Front Oncol 12, 1118675 (2023).

20. Chakrabarty, R. P. & Chandel, N. S. Mitochondria as Signaling Organelles Control Mammalian Stem Cell Fate. Cell Stem Cell 28, 394–408 (2021).

21. Losman, J. A., Koivunen, P. & Kaelin, W. G. 2-Oxoglutarate-dependent dioxygenases in cancer. Nat Rev Cancer 20, 710–726 (2020).

22. Li, X. et al. Inhibition of fatty acid oxidation enables heart regeneration in adult mice. Nature 622, 619–626 (2023).

23. Chen, Y. et al. Reversible reprogramming of cardiomyocytes to a fetal state drives heart regeneration in mice. Science (1979) 373, 1537–1540 (2021).

24. Chen, S. et al. The role of glycolytic metabolic pathways in cardiovascular disease and potential therapeutic approaches. Basic Res Cardiol 118, 48 (2023).

25. Morrisey, E. E. Rewind to recover: Dedifferentiation after cardiac injury. Cell Stem Cell 9, 387– 388 (2011).

26. Liu, W. et al. Medium acidosis drives cardiac differentiation during mesendoderm cell fate specification from human pluripotent stem cells. Stem Cell Reports 19, 1304–1319 (2024).

27. Arnold, Z. et al. Tenascin-C drives cardiovascular dysfunction in a mouse model of diabetic cardiomyopathy. Cardiovasc Diabetol 24, (2025).

28. Ehler, E., Moore-Morris, T. & Lange, S. Isolation and culture of neonatal mouse cardiomyocytes. Journal of Visualized Experiments 50154 (2013) doi:10.3791/50154.

29. Ackers-Johnson, M. et al. A Simplified, Langendorff-Free Method for Concomitant Isolation of Viable Cardiac Myocytes and Nonmyocytes From the Adult Mouse Heart. Circ Res 119, 909–920 (2016).

30. Badstöber, J., Gachon, C. M. M., Ludwig-Müller, J., Sandbichler, A. M. & Neuhauser, S. Demystifying biotrophs: FISHing for mRNAs to decipher plant and algal pathogen-host interaction at the single cell level. Sci Rep 10, (2020).

31. Snaebjornsson, M. T. et al. Targeting aldolase A in hepatocellular carcinoma leads to imbalanced glycolysis and energy stress due to uncontrolled FBP accumulation. Nat Metab 7, 348–366 (2025).

32. Pircher, H. et al. Identification of Human Fumarylacetoacetate Hydrolase Domain-containing Protein 1 (FAHD1) as a Novel Mitochondrial Acylpyruvase. Journal of Biological Chemistry 286, 36500–36508 (2011).

