## Supplemental Table 1 for "A novel role for Oxaloacetate Decarboxylase FAHD1 in cardiomyocyte maturation"

| Metabolite | LS23-026_KO_1 | LS23-026_KO_2 | LS23-026_KO_3 |
|---|---|---|---|
| Group | 2 - KO | 2 - KO | 2 - KO |
| 2-Hydroxyglutaric acid6.999neg | 9.311321902 | 6.047949797 | 8.825037847 |
| Oxoglutaric acid6.166neg | 12.08037518 | 10.62288448 | 10.96018693 |
| Citric acid7.592neg | 127.2180596 | 75.53864712 | 110.7898327 |
| Fumaric acid6.379neg | 547.4210845 | 500.6318485 | 412.8740791 |
| Glycine6.762neg | 80.19738057 | 98.21888877 | 78.72951057 |
| L-Lactic acid5.414neg | 2385.268409 | 2012.745505 | 1735.001354 |
| L-Asparagine6.983neg | 25.82197804 | 23.46240075 | 23.50155722 |
| L-Aspartic acid6.715neg | 202.6562702 | 174.6131424 | 232.0006343 |
| L-Glutamic acid6.545neg | 553.1952975 | 567.8175375 | 563.9115939 |
| L-Glutamine6.725neg | 817.2002172 | 641.0695356 | 692.5530617 |
| L-Histidine10.869neg | 34.16226731 | 32.38135327 | 37.22442096 |
| L-Lysine12.349neg | 69.1012236 | 74.10597632 | 65.19794565 |
| L-Valine6.146neg | 34.03129528 | 46.54753077 | 29.35297222 |
| Malic acid6.561neg | 252.9030598 | 129.0569013 | 157.7338721 |
| Pyruvic acid1.743neg | 6681.557642 | 4494.019474 | 2018.758649 |
| Succinic acid6.248neg | 23.81418541 | 18.75606187 | 22.61884895 |
| Creatine6.512pos | 2566.882245 | 2027.685438 | 2336.840724 |
| L-Alanine6.57pos | 938.3968313 | 894.3911675 | 846.3786566 |
| L-Arginine11.969pos | 38.67163435 | 39.88274573 | 29.55413781 |
| L-Isoleucine5.844pos | 25.1111752 | 35.5565405 | 24.02039702 |
| L-Methionine6.023pos | 15.29322372 | 22.08737502 | 14.07208956 |
| L-Phenylalanine5.47pos | 159.2966572 | 288.3750724 | 173.0522921 |
| L-Proline6.355pos | 10.1442076 | 18.93854254 | 18.52964042 |
| L-Serine6.847pos | 0.946461197 | 0.660228046 | 0.553715429 |
| L-Threonine6.716pos | 78.07583403 | 60.97339886 | 73.14919359 |
| L-Tryptophan5.419pos | 68.44901978 | 86.833734 | 28.8392129 |
| L-Tyrosine6.03pos | 0.396572848 | 0.702100409 | 0.275605891 |
| Ornithine11.968pos | 5.628589438 | 10.00941454 | 5.104269798 |
| 3-Hydroxybutyric acid4.338neg | 13882124.72 | 12251427.13 | 15980870.02 |
| 3-Phosphoglyceric acid7.034neg | 4179889.685 | 3253700.414 | 2396432.736 |
| ADP7.16neg | 6767483.651 | 2962992.811 | 2268198.443 |
| AMP6.703neg | 157630943.1 | 146368250.7 | 131096945.7 |
| Ascorbic acid6.356neg | 7121405.464 | 5390323.294 | 5466180.034 |
| ATP7.101neg | 788558.9326 | 101695.7418 | 169181.014 |
| CDP7.02neg | 19054.53795 | 13691.82513 | 11109.57254 |
| cis-Aconitic acid6.327neg | 16427804.11 | 11181278.03 | 17182988.21 |
| CMP6.765neg | 987140.611 | 765920.6826 | 605217.0284 |
| CTP7.097neg | 30401.76443 | 0 | 0 |
| Dihydroxyacetone phosphate6.865neg | 716066.7153 | 551743.9014 | 417946.605 |
| Fructose4.736neg | 975477.5982 | 684564.5394 | 630046.6883 |
| Fructose16-bisphosphate7.402neg | 125870.0126 | 96168.02059 | 51879.05958 |
| GDP7.161neg | 78850.15141 | 42574.71826 | 30328.56037 |
| Glucosamine 6-phosphate5.625neg | 0 | 0 | 0 |
| Glucose5.875neg | 5138927.092 | 8267917.778 | 4687965.446 |

| | | | |
|---|---|---|---|
| Glucose-6-Phosphate6.895neg | 6594860.241 | 5313558.026 | 1704432.536 |
| Glutathione6.765neg | 15706147.27 | 11943945.41 | 10587726.99 |
| Glyceric acid6.187neg | 4552291.074 | 3476817.826 | 6698340.719 |
| Glycerol-3-phosphate6.76neg | 66767883.84 | 44191687.23 | 40492321.97 |
| GMP6.803neg | 6367549.359 | 6084514.544 | 4959267.697 |
| GTP7.092neg | 49110.82138 | 15640.22352 | 7581.194474 |
| Hypoxanthine2.249neg | 42396020.69 | 64838884.63 | 61614506.98 |
| IMP6.67neg | 29475507.89 | 15169252 | 13604562.11 |
| L-Cysteine7.349neg | 0 | 15144.23904 | 0 |
| L-Cystine7.448neg | 21944.73145 | 69771.8201 | 18035.63095 |
| myo-Inositol6.714neg | 4666282.919 | 3039958.174 | 2940019.41 |
| N-Acetyl-L-aspartic Acid6.303neg | 18934926.76 | 44671213.93 | 44600295.16 |
| Oxidized glutathione7.076neg | 7805058.967 | 7462118.652 | 4702736.641 |
| pa_Folate6.371neg | 0 | 0 | 0 |
| pa_Itaconic acid6.338neg | 6204696.137 | 4874942.715 | 7053626.49 |
| pa_Sedoheptulose 7-phosphate6.766neg | 430057.2188 | 490767.4154 | 185138.408 |
| Pantothenic acid5.462neg | 57329447.38 | 53316109.92 | 63427302.46 |
| Phosphoserine7.054neg | 0 | 0 | 0 |
| Ribose-5-Phosphate6.602neg | 14037316.22 | 9905460.114 | 6564660.974 |
| UDP6.807neg | 412620.9606 | 233227.9639 | 263000.2109 |
| UDP-glucoronic acid6.664neg | 323775.4333 | 200311.4764 | 368489.7057 |
| UDP-glucose6.512neg | 1246985.425 | 883894.3528 | 1121420.141 |
| UDP-N-acetylglucosamine6.417neg | 1867987.816 | 1400890.969 | 1375271.286 |
| UMP6.69neg | 5022436.472 | 3879384.277 | 2924842.509 |
| Uridine2.143neg | 16999304.1 | 19265421.43 | 26392407.39 |
| UTP7.028neg | 403482.5418 | 72636.80521 | 92395.43696 |
| Xanthine2.883neg | 32069188.81 | 30637393.03 | 33803123.02 |
| 4-Hydroxyproline6.686pos | 80118.90168 | 75577.42823 | 363139.4666 |
| 5-Oxoproline6.731pos | 52494407.55 | 38270207.6 | 44329288.76 |
| Acetylcarnitine6.633pos | 13253743.01 | 20438636.56 | 29834700.81 |
| Adenine1.933pos | 88839395.63 | 89734263.15 | 149536955 |
| Adenosine2.284pos | 587004308.8 | 580467849.8 | 947967856.9 |
| Betaine6.138pos | 331677288.4 | 358840075.6 | 409569372.5 |
| Butanoyl-Carnitine5.937pos | 22974478.47 | 12319313.95 | 72337981.26 |
| CDP-choline6.893pos | 2920740.988 | 1837035.447 | 2172809.256 |
| Choline10.593pos | 152038533.8 | 124699597.9 | 139303452.2 |
| Citrulline6.892pos | 3713335.593 | 2954349.141 | 3339855.34 |
| Creatinine3.169pos | 169640856.5 | 164717027.7 | 203453111.3 |
| Cytidine5.001pos | 9999619.046 | 7394749.253 | 9706537.947 |
| Cytosine3.644pos | 3270452.042 | 4190944.582 | 5181730.376 |
| Cytosine5.061pos | 21183554.04 | 16715370.15 | 22609637.24 |
| FAD6.235pos | 1223633.876 | 780872.1208 | 1120740.676 |
| gamma-Aminobutyric acid6.887pos | 562056.0947 | 530883.5314 | 433010.1322 |
| gamma-Glutamylleucine6.251pos | 161512.731 | 150135.5258 | 91104.45504 |
| gamma-Glutamylvaline6.353pos | 145747.8083 | 197726.6904 | 103517.8021 |
| gamma-L-Glutamyl-L-alanyl-glycine6.65pos | 226089.5882 | 105432.264 | 58247.85381 |

| | | | |
|---|---|---|---|
| Glucosamine6.054pos | 9731.999615 | 10683.25669 | 7933.02595 |
| Inosine3.972pos | 35152814.86 | 32071434.79 | 42191649.82 |
| L-Carnitine7.09pos | 859458706.5 | 901769533.1 | 1159948101 |
| N2-Acetylornithine6.959pos | 65764.69895 | 109822.0006 | 108192.0013 |
| NAD6.618pos | 13720478.22 | 8221151.167 | 12248121.92 |
| NADH6.61pos | 563908.0733 | 462350.1002 | 319102.8255 |
| NADP6.901pos | 543729.8869 | 289294.6586 | 333796.2249 |
| pa_gamma-Glutamylalanine6.573pos | 113686.3272 | 34017.98663 | 33042.04264 |
| pa_gamma-Glutamylthreonine6.665pos | 41867.36845 | 0 | 0 |
| pa_Histamine12.049pos | 8753412.647 | 11850759.9 | 6170673.259 |
| pa_Spermidine5.803pos | 2445128.262 | 4823901.599 | 9691505.85 |
| pa_Spermine5.912pos | 2047882.814 | 5889974.446 | 7658254.429 |
| p-Aminobenzoic acid6.259pos | 12940137.43 | 19653261.75 | 27078022.58 |
| Phosphocreatine6.668pos | 67870.2958 | 0 | 0 |
| Propionyl-Carnitine6.274pos | 282310961.3 | 206413725.3 | 221648927.8 |
| Pyridoxine1.76pos | 15321.11779 | 286509.2109 | 26456.70025 |
| Riboflavin3.416pos | 55485.6604 | 111273.1625 | 100642.0831 |
| S-Adenosylmethionine11.777pos | 1932439.766 | 1241702.792 | 1187181.691 |
| Taurine5.955pos | 330858093.9 | 306570935.3 | 418960285.6 |
| Thiamine11.846pos | 26485.78987 | 44435.44218 | 32325.72451 |
| Thymine6.267pos | 16279.00803 | 0 | 0 |
| Uracil2.245pos | 2194381.142 | 2786439.622 | 3975227.297 |

| LS23-026_KO_4 | LS23-026_WT_1 | LS23-026_WT_2 | LS23-026_WT_3 | LS23-026_WT_4 |
|---|---|---|---|---|
| 2 - KO | 1 - WT | 1 - WT | 1 - WT | 1 - WT |
| 14.28263216 | 7.345009148 | 9.83806774 | 10.10680941 | 9.979454932 |
| 11.3656305 | 8.195799607 | 10.78856201 | 16.20568207 | 14.54835396 |
| 99.46019619 | 53.28423467 | 90.06842044 | 124.6235841 | 111.8243402 |
| 554.5859357 | 522.0289656 | 203.8760536 | 446.9796111 | 278.5868135 |
| 97.05817931 | 61.21336957 | 63.38614506 | 86.64340647 | 73.79097214 |
| 3048.138866 | 1992.559265 | 2117.53268 | 2509.225184 | 2479.40667 |
| 28.13961467 | 16.6802246 | 22.34600607 | 22.21481579 | 17.61062805 |
| 179.5207109 | 119.1355495 | 173.7750725 | 235.2260688 | 163.7619066 |
| 715.191928 | 404.1537668 | 509.7818083 | 574.099686 | 544.9159449 |
| 754.897632 | 585.0016729 | 723.7063598 | 825.1079871 | 785.1832083 |
| 44.06267103 | 27.2920527 | 27.12514584 | 31.53675119 | 28.23811407 |
| 72.90809614 | 61.25874893 | 60.44565279 | 61.878428 | 64.54278349 |
| 52.12236373 | 38.26662719 | 30.70118962 | 35.72447281 | 34.67042643 |
| 183.3006739 | 205.3950423 | 149.4365341 | 272.8863474 | 145.5370921 |
| 5143.249233 | 29706.03218 | 4694.220964 | 66623.23557 | 17006.56438 |
| 26.02191958 | 21.54161098 | 47.32466923 | 38.70787385 | 38.35185672 |
| 2763.555398 | 1717.66197 | 2007.500268 | 2676.809959 | 2008.176051 |
| 1043.732201 | 911.6780584 | 1050.527709 | 1216.274814 | 1174.086906 |
| 31.88787935 | 31.56161563 | 43.6055177 | 41.20070129 | 37.93961394 |
| 47.74703149 | 23.76573464 | 21.75911236 | 25.46602453 | 24.61806542 |
| 23.93068173 | 38.03386222 | 17.4257623 | 28.16539907 | 31.94092534 |
| 345.1950759 | 111.43677 | 148.4944342 | 137.7546492 | 154.9500169 |
| 22.51587531 | 15.09614134 | 12.05746797 | 12.49616779 | 15.26506502 |
| 0.762695845 | 0.821312144 | 0.618805922 | 0.903775514 | 0.702544248 |
| 85.42457607 | 75.80794685 | 61.0373859 | 78.50265208 | 73.68269811 |
| 73.70786868 | 85.72613114 | 84.76911452 | 58.14869004 | 62.13053792 |
| 0.309228248 | 1.380202882 | 0.315195902 | 0.520228453 | 0.274232652 |
| 11.04569957 | 8.106492428 | 6.21137175 | 5.462528842 | 7.69975118 |
| 23234168.92 | 3669479.417 | 13261197.33 | 8995184.121 | 19654551.71 |
| 2354977.484 | 2757093.838 | 4643250.911 | 5572572.1 | 4690778.663 |
| 913759.2778 | 5571574.164 | 5641450.062 | 9187868.029 | 4965984.736 |
| 132404428 | 62594404.31 | 111698809.8 | 174938060.4 | 117417193.8 |
| 5502575.878 | 5535035.606 | 6018404.734 | 9551886.509 | 6425870.894 |
| 24869.94011 | 2898633.436 | 427234.9923 | 291924.8982 | 215555.9632 |
| 18818.78515 | 157216.9913 | 24252.12259 | 15017.19636 | 0 |
| 13419200.5 | 5304622.523 | 10577549.89 | 15365050 | 18040873.9 |
| 643499.5106 | 669291.4359 | 667890.9381 | 856153.626 | 837298.66 |
| 0 | 147289.4827 | 11191.97551 | 0 | 12937.79137 |
| 825678.0006 | 576779.039 | 682808.2885 | 1302203.906 | 965158.867 |
| 1135136.927 | 775476.1247 | 687007.6886 | 823364.7006 | 937911.1926 |
| 93849.58654 | 150822.9118 | 137342.4288 | 329054.8437 | 131613.1945 |
| 19669.38187 | 122877.8183 | 53090.72011 | 80903.80478 | 91375.39747 |
| 0 | 0 | 0 | 0 | 0 |
| 6268853.674 | 13257124.98 | 6053398.699 | 10555854.1 | 7509175.132 |

| | | | | |
|---|---|---|---|---|
| 28058726.57 | 16983434.25 | 15077573.66 | 12174844.73 | 4994881.69 |
| 10261881.92 | 11311377.5 | 11534087.58 | 13452716.4 | 9730048.205 |
| 4347613.43 | 1971920.368 | 3815518.244 | 4141644.339 | 5297397.405 |
| 57572191.73 | 50598563.82 | 53673349.47 | 51225166.51 | 52685862.17 |
| 5310900.512 | 5538569.811 | 4310511.909 | 4938916.533 | 3979861.24 |
| 0 | 75051.55649 | 26254.05934 | 21153.30924 | 12178.69591 |
| 91360637.21 | 37044674.23 | 40112285.25 | 30957785.23 | 58122349.91 |
| 13450172.16 | 59423467.25 | 28840004.84 | 29556555.44 | 17120839.19 |
| 0 | 9650.576382 | 0 | 10700.12885 | 0 |
| 58645.69882 | 50760.25407 | 12763.81441 | 73421.45412 | 25165.07811 |
| 3318744.216 | 2938113.889 | 2664334.844 | 4385098.668 | 4271641.034 |
| 82450521.01 | 15434391.56 | 19098443.69 | 33657767.59 | 47439820.45 |
| 3488658.667 | 7640972.266 | 6370210.077 | 6857471.725 | 7447981.44 |
| 0 | 0 | 0 | 0 | 0 |
| 7367215.666 | 2483741.849 | 4544543.345 | 6535978.737 | 7419237.921 |
| 410253.2137 | 518325.085 | 290363.7671 | 328306.2325 | 290032.5212 |
| 92705010.68 | 41402322.24 | 43554583.31 | 63323237.2 | 49156827.01 |
| 0 | 0 | 0 | 0 | 0 |
| 9239145.144 | 14252693.12 | 8204466.657 | 7354750.869 | 7306566.887 |
| 374941.7519 | 1040254.565 | 288893.2816 | 434671.755 | 334852.8907 |
| 97820.34265 | 475043.7769 | 183265.493 | 197775.489 | 279580.1857 |
| 354201.009 | 716283.3022 | 718549.8177 | 665585.478 | 1163579.949 |
| 1409075.917 | 1223029.761 | 1535732.299 | 1898329.794 | 1548411.13 |
| 5018465.567 | 3988114.728 | 4892489.233 | 6689081.222 | 3867311.259 |
| 24684339.41 | 10574825.46 | 12440301.77 | 14630331.95 | 18234121.08 |
| 0 | 1484411.474 | 180348.9501 | 0 | 96005.45303 |
| 54193096.75 | 15331000.43 | 21392387.92 | 21693965.89 | 30561077.12 |
| 343598.2002 | 67335.16911 | 46483.54521 | 70909.2718 | 49862.75882 |
| 50211369.5 | 32996129.46 | 43662877.4 | 46602789.48 | 40224785.19 |
| 33205297.2 | 12407808.48 | 19638392.12 | 18847642.17 | 25052757.73 |
| 124808051.4 | 22210661.29 | 55761508.43 | 70293709.13 | 69074378.95 |
| 783000535.6 | 138613997.5 | 343103614.1 | 410857578.2 | 417321206.5 |
| 528736205.5 | 221445926.7 | 318129374 | 383685352.7 | 436420641.4 |
| 15120915.53 | 5974189.449 | 40870484.71 | 21772840.91 | 41318377.18 |
| 2453037.508 | 1607824.722 | 2305386.851 | 2906760.682 | 1949711.488 |
| 155665968.4 | 119680011.1 | 103551959.6 | 117524544.3 | 114207042.6 |
| 3419852.469 | 2299632.386 | 2394978.622 | 3386636.284 | 2881202.81 |
| 252944758.1 | 79996640.35 | 163042568.3 | 174913985.9 | 159450385.7 |
| 7843724.922 | 4745529.633 | 6922341.728 | 5305125.617 | 6482996.687 |
| 6163113.273 | 2365261.947 | 3505480.005 | 4076640.516 | 4002226.192 |
| 17653935.5 | 10718876.14 | 16128708.23 | 11909259.9 | 14571554.66 |
| 542155.0732 | 600347.1242 | 1057198.997 | 1245404.035 | 892687.8274 |
| 693894.4801 | 278672.3057 | 472148.7556 | 523720.7344 | 630835.8278 |
| 178506.6094 | 172647.2734 | 166943.7451 | 133137.2195 | 147671.0061 |
| 123312.5069 | 151563.6787 | 130454.9703 | 91611.86914 | 158629.6709 |
| 46862.72281 | 61603.39161 | 100685.9911 | 205034.5838 | 195734.9658 |

| | | | | |
|---|---|---|---|---|
| 0 | 21899.95552 | 3513.844062 | 17702.71339 | 10688.14365 |
| 49426468.62 | 20639650.39 | 27260311.15 | 25077849.52 | 34047062.26 |
| 1455387749 | 520770249.3 | 860078357.8 | 960781105.4 | 1018008207 |
| 217909.3021 | 73511.79794 | 86344.44151 | 72671.18015 | 72839.34264 |
| 7939179.341 | 3146230.252 | 12370165.15 | 17386433.57 | 15728205.18 |
| 392058.4967 | 122903.5719 | 613534.1398 | 635059.9514 | 569605.5849 |
| 72322.11107 | 283913.8917 | 380788.4454 | 553175.012 | 348447.8046 |
| 75710.75612 | 73039.94736 | 104907.0207 | 40896.02613 | 39492.56925 |
| 0 | 0 | 0 | 0 | 0 |
| 7291031.372 | 5465055.621 | 4928591.616 | 8887722.765 | 7202302.626 |
| 16873097.81 | 1747937.759 | 2752037.909 | 4889548.763 | 11840565.5 |
| 5945845.868 | 908343.7557 | 2796502.123 | 3660821.463 | 4680846.534 |
| 49576692.83 | 8760186.558 | 17625904.98 | 17533903.19 | 20666946.1 |
| 0 | 0 | 0 | 32560.47244 | 164335.5821 |
| 400080954 | 143035943.5 | 180241629.8 | 267021593 | 224094537.2 |
| 98797.79886 | 216654.0501 | 29645.3864 | 19666.82484 | 137540.6313 |
| 142620.0215 | 35767.41916 | 38999.61742 | 40464.30787 | 32168.28991 |
| 1464608.504 | 816440.5376 | 1849927.604 | 2067442.634 | 1390584.089 |
| 488128974 | 183047184 | 298879110.7 | 373994670.9 | 340263948.8 |
| 41992.7396 | 32539.44318 | 33778.99525 | 26621.26142 | 29503.50173 |
| 26752.08173 | 11117.06149 | 14166.35228 | 0 | 19126.83238 |
| 3740915.915 | 1478989.483 | 2219966.895 | 2263668.199 | 2366446.849 |
